# Host prion protein expression levels impact prion tropism for the spleen

**DOI:** 10.1101/2019.12.17.879676

**Authors:** Vincent Béringue, Philippe Tixador, Olivier Andréoletti, Fabienne Reine, Johan Castille, Thanh-Lan Laï, Annick Le Dur, Aude Laisné, Laetitia Herzog, Bruno Passet, Human Rezaei, Jean-Luc Vilotte, Hubert Laude

**Affiliations:** VIM, INRA, Université Paris-Saclay, Jouy-en-Josas, France; IHAP, INRA, ENVT, Toulouse, France; GABI, INRA, AgroParisTech, Université Paris-Saclay, Jouy-en-Josas, France

## Abstract

Prions are pathogens formed from abnormal conformers (PrP^Sc^) of the host-encoded cellular prion protein (PrP^C^). PrP^Sc^ conformation to disease phenotype relationships extensively vary among prion strains. In particular, prions exhibit a strain-specific tropism for lymphoid tissues. Prions can be composed of several substrain components. There is evidence that these substrains can propagate in distinct tissues (e.g. brain and spleen) of a single individual, providing an experimental paradigm to study the cause of prion tissue selectivity. Previously, we showed that PrP^C^ expression levels govern prion substrain selection in the brain. Transmission of sheep scrapie isolates (termed LAN) to multiple lines of transgenic mice expressing varying levels of ovine PrP^C^ in the brain resulted in the phenotypic expression of the dominant sheep substrain in mice expressing near physiological PrP^C^ levels, whereas a minor substrain replicated preferentially on high-expressors. Considering that PrP^C^ expression levels are markedly decreased in the spleen compared to the brain, we interrogate whether spleen PrP^C^ dosage could drive prion selectivity. The outcome of the transmission of a large cohort of LAN-like scrapie isolates in the spleen from high-expressors correlated with the replication rate dependency on PrP^C^ amount; There was a prominent spleen colonization by the substrain preferentially replicating on low-expressors and a relative incapacity of the substrain with higher-PrP^C^ level need to propagate in the spleen. Early colonization of the spleen allowed neuropathological expression of the lymphoid substrain after intraperitoneal inoculation. In addition, a pair of substrain variants resulting from the adaptation of human prions to ovine high-expressors, and exhibiting differing brain versus spleen tropism, showed different tropism on transmission to low-expressors, with the lymphoid substrain colonizing the brain. Overall, these data suggest that PrP^C^ expression levels are instrumental in prion substrain lymphotropism.

**Author summary:** The cause of prion phenotype variation among prion strains remains poorly understood. In particular prions replicate in a strain-dependent manner in the spleen. This can result in prion asymptomatic carriers. Based on our previous observations that dosage of the prion precursor (PrP) determined prion substrain selection in the brain, we examine whether PrP levels in the spleen could drive prion replication in this tissue, due to the low levels of the protein. We observe that the prion substrain with higher PrP need for replication did barely replicate in the spleen, while the component with low PrP need replicated efficiently. In addition, other human co-propagating prions with differing spleen and brain tropism showed different tropism on transmission to mice expressing low PrP levels, with the lymphoid substrain colonizing the brain. PrP^C^ expression levels may thus be instrumental in prion tropism for the lymphoid tissue. From a diagnostic point of view, given the apparent complexity of prion diseases with respect to prion substrain composition, these data advocate to type extraneural tissues or fluids for a comprehensive identification of the circulating prions in susceptible mammals.

## Introduction

Mammalian prions are proteinaceous pathogens causing fatal neurodegenerative diseases termed transmissible spongiform encephalopathies (TSE) in humans and animals. TSE include scrapie in sheep and goats, bovine spongiform encephalopathy (BSE) in cattle, chronic wasting disease (CWD) in cervid populations and Creutzfeldt-Jakob disease (CJD) in humans [1]. Prions are formed from abnormal, β-sheet enriched conformers (PrP^Sc^) of the host encoded cellular prion protein (PrP^C^). Prions replicate by templating the conversion and polymerization of PrP^C^ by an autocatalytic process [2, 3]. Multiple strains of prions are recognized phenotypically within the same host species. Strains are conformational variants of PrP^Sc^, at the level of the tertiary and/or quaternary structure [4–6]. In the infected host, prion strains exhibit specific incubation periods, stereotyped clinical signs and neuropathology, and specific tropism for the central nervous system (CNS) and lymphoid tissues (for review [7, 8]). Thus, certain prions can replicate early and at fairly high levels in tissues of the lympho-reticular system such as the Peyer’s patches, spleen and lymph nodes [9–11]. Lymphotropic prions are then transported from these early reservoirs of infectivity to the brain either by the enteric nervous system (Peyer’s patches) or by the peripheral nervous system and the spinal cord (spleen, lymph nodes) [12–14]. After peripheral infection, the presence of differentiated follicular dendritic cells (FDC) is required for efficient prion replication in the lymphoid tissue and subsequent neuroinvasion [9, 15–21]. Yet, prions can differ in their capacity to neuroinvade. Certain prions can persist in the lymphoid tissue without accessing the CNS [22–25]. Of particular concern are BSE/variant CJD prions, which are suspected to accumulate in human lymphoid tissue without neuroinvasion. 1:2000 exposed individuals in the UK may be silent carriers in the lymphoid tissue, causing risks of secondary transmission [26].

Although one dominant PrP^Sc^ conformation is usually detected by conventional immunodetection methods, there is clear evidence that natural or experimental prion sources can be composed of several substrains, in variable proportions, and with differing PrP^Sc^ conformations [27–30]. As a result, experimental prion transmission, in a homotypic context (i.e. host PrP^C^ and prion PrP^Sc^ share identical primary sequence) or not, can lead to the isolation of different substrains in the brain and spleen of a single transgenically modified mouse expressing PrP [22, 23, 28, 31]. The reasons for such substrain segregation between the brain and the spleen and more generally for the incapacity of certain strains to replicate in the spleen remain poorly understood. It has been proposed that tertiary or quaternary PrP^Sc^ conformations impact peripheral prion replication [25, 31–33]. The aforementioned tissue segregative transmission provides a relevant experimental paradigm to study prion lymphotropism.

Intracerebral transmission of natural sheep scrapie isolates (referred to as LAN isolates [9, 34]) to multiple lines of transgenic mice expressing ovine PrP^C^ (VRQ allele at codons 136, 154 and 171 of the PrP-encoding gene, where V, R, and Q stand for valine, arginine, and glutamine, respectively) and at variable levels revealed that the PrP^C^ expression level in the brain critically determines prion substrain selection [29]. The so-called LA21K dominant component in the sheep brain (giving the strain signature of LAN isolates) was selected in the brain of mice expressing near physiological PrP^C^ levels, while preferential selection of the so-called LA19K prion subcomponent occurred in the brain of ‘high-expressor’ mice.

PrP^C^ levels are markedly lowered in the spleen, - approximately 20-fold compared to the brain-, in wild-type mice as in some high-expressor transgenic mouse models [22]. We thus interrogate whether PrP^C^ expression levels could impact prion peripheralization. We compare the outcome of the transmission of a large cohort of sheep scrapie isolates, including the LAN-like isolates, in the brain and spleen tissue of high-expressors. In accordance with their affinity for PrP^C^ levels, we show preferential replication of LA21K and absence of replication of LA19K prions in the spleen. Consequently, varying the inoculation route allows the dominant expression of LA21K prions in the brain of high-expressors. Further, we show that a pair of strains resulting from the adaptation of cortical MM2 CJD subtype to ovine high-expressors, and exhibiting differing brain versus spleen tropism (T1^Ov^ strain in spleen and T2^Ov^ strain in brain [28]) showed opposite tropism in low-expressors. Collectively, these data raise the possibility that PrP^C^ expression levels are instrumental in specific tropism for the lymphoid tissue.

## Methods

### Ethics statement

Animal care and experiments were conducted in strict compliance with ECC and EU directives 86/009 and 2010/63. They were reviewed and approved by the local ethics committee of the author’s institution, (name: COMETHEA: Comité d’Ethique en Expérimentation Animale du Centre INRA de Jouy-en-Josas et AgroParisTech). The permit numbers delivered by the COMETHEA are 12/034 and 15/056.

### Transgenic mice

As high-expressors, we used tg338 mice that overexpress the VRQ allele of ovine PrP. The transgene construct consists of a large DNA sequence derived from ovine BAC libraries, encompassing natural regulatory sequences of the PrP gene transcription unit [35]. The PrP^C^ levels in the brain are ∼8-fold higher than in the sheep brain [29]. The spleen-to-brain PrP^C^ ratio (∼1:20) in tg338 mice is comparable to that found in conventional mouse models [22]. The strongest PrP^C^ staining in tg338 spleens is FDC-associated [22]. Thus, there is no aberrant expression of PrP^C^ in the spleen of these mice, quantitatively or qualitatively. As low-expressor mice, we used tg335^+/-^ (same transgene construct as the tg338 mice) and tg143^+/-^ mice, which express 1.2-fold and 1.5-fold PrP^C^ as compared to sheep brain, respectively [29, 35].

### TSE sources

Brain and spleen from sheep terminally affected with natural scrapie (LAN404 isolate) were provided by the Institut National de la Recherche Agronomique (O. Andréoletti, Toulouse, France). Brain extracts from French, Dutch, Spanish, Irish, and English sheep scrapie isolates were provided by the Institut National de la Recherche Agronomique (O. Andréoletti, Toulouse, France), the Central Veterinary Institute (J.M. Langeveld, Wageningen, Lelystad, Netherlands), the French National TSE Reference Laboratory (T. Baron, Anses, Lyon, France), CISA-INIA (J.M. Torres, Madrid, Spain), the Central Veterinary Research Laboratory, (E. Monks, Dublin, Ireland) and the former European TSE Reference Laboratory (J. Spiropoulos, VLA, Addlestone, UK). The CH1641 sheep scrapie strain [36] was provided by the Institute for Animal Health (N. Hunter, Edinburgh, UK). The TSE isolates are detailed in supplemental Table 1.

The T1^Ov^ and T2^Ov^ strains were isolated after serial transmission of a human sporadic CJD brain (MM2, rare cortical form) to ovine PrP tg338 mice [28]. As inoculum, a pool of brains from tg338 mice at the 4^th^ serial passage of MM2-CJD and containing T1^Ov^ and T2^Ov^ prions, with T2^Ov^ in higher proportion, was used. For comparison, pools of brains from tg338 mice inoculated with cloned T1^Ov^ or bicloned T2^Ov^ prions were used [28].

### Mouse transmission assays

Sheep tissue extracts were prepared as 10% w/v homogenate in 5% w/v glucose with a Precellys rybolyzer (Ozyme, Montigny-le-Bretonneux, France). To avoid any cross-contamination, a strict protocol based on the use of disposable equipment and preparation of all inocula in a class II microbiological cabinet was followed. For intracerebral inoculations, twenty microliters were inoculated in the right hemisphere to groups of individually identified mice, at the level of the parietal cortex. For intraperitoneal inoculations, hundred microliters of a 2% w/v solution in 5% glucose were used. For subsequent passage, mouse brains and spleens were collected with dedicated, sterile, disposable tools, homogenized at 20% w/v in 5% glucose; twenty microliters were reinoculated intracerebrally at 10% w/v. Animals were supervised daily for TSE development. Animals at terminal stage of disease or at end life were euthanized. To study the kinetics of PrP^Sc^ accumulation in the spleen and in the brain after intraperitoneal or intracerebral infection, mice were euthanized healthy in triplicates at regular time-points post-inoculation, as indicated. For immunoblot analyses, brains and spleens were immediately frozen at -80°C until use. For histoblot analyses, the collected brains were frozen on dry ice before storage at -80°C.

### Immunoblot analyses

Brains and spleens were analyzed for proteinase K (PK)-resistant PrP^Sc^ (PrP^res^) content using a previously published protocol [28]. Briefly, PrP^res^ was extracted from 20 % w/v tissue homogenates with the Bio-Rad TeSeE detection kit. Aliquots were digested with PK (200 μg/ml final concentration) for 10 min at 37 °C before B buffer precipitation and centrifugation at 28,000 × g for 15 min. Pellets were resuspended in Laemmli sample buffer, denatured, run on 12 % Bis/Tris gels (Bio-Rad), electrotransferred onto nitrocellulose membranes, and probed with 0.1 μg/ml biotinylated anti-PrP monoclonal antibody Sha31 antibody (human PrP epitope 145-152, [37]) or with 0.1 μg/ml anti-PrP 12B2 antibody (human PrP epitope 89-93, epitope, [38]) and followed by streptavidin conjugated to horseradish peroxidase (HRP) or by HRP conjugated to goat anti-mouse IgG1 antibody (1/20 000 final dilution), respectively. Immunoreactivity was visualized by chemiluminescence (GE Healthcare). The relative amounts of PrP^res^ glycoforms were determined by the use of GeneTools software after acquisition of chemiluminescent signals with a GeneGnome digital imager (Syngene, Frederick, MD).

### Histoblot analyses

Brain cryosections were cut at 8-10 μm, transferred onto Superfrost slides and kept at -20 °C until use. Histoblot analyses were performed as described [39], using the 12F10 anti-PrP antibody (human PrP epitope 142-160, [40]). Analysis was performed with a digital camera (Coolsnap, Photometrics) mounted on a binocular glass (SZX12, Olympus). The sections presented are representative of the analysis of three brains samples.

### Resistance to proteinase K digestion

20% (w/v) brain homogenates from tg338 mice infected with 127S and LA19K prions were diluted at 5% in a solubilization buffer (20 mM HEPES pH 7.4, 150 mM NaCl, 5 mM EDTA, 1 mM DTT, 2% (w/v) dodecyl-β-D-maltoside (Sigma)) 2% N-lauryl sarcosine (Fluka), final concentrations) and digested for 2h at 37°C with increasing concentrations of PK (0 to 10 000 μg/ml), as indicated. After digestion, the samples were diluted in equal volumes of Laemmli buffer, denatured and analyzed for PrP content by western blot, as above.

### PrP^Sc^ degradation by primary cultured peritoneal macrophages

To induce the multiplication of peritoneal macrophages, healthy tg338 mice were intraperitoneally injected with 3% Brewer thioglycolate broth (BD Biosciences). Three days after the injection, mice were euthanized by cervical column disruption. Peritoneal lavage was performed with 5 ml D-PBS (Gibco). The lavage fluid was mixed with an equivalent volume of 4°C Dulbecco’s Modified Eagle Medium (DMEM, Lonza) supplemented with 10% fetal calf serum (FCS, Biowhitaker), streptomycin and penicillin (PS, Gibco) before centrifugation at 100 x g for 5 min. Red cells were lyzed with hematolytic medium (155 mM ammonium chloride, 12 mM sodium carbonate, pH 7.4). The cells were resuspended in DMEM-10% FCS-PS and washed twice by centrifugation. After a live cell count, the cells were aliquoted in 6-well plates (6.10^6^ cells / well) and incubated at 37°C. After 24h, non-adhering cells were discarded by washing the wells with D-PBS. Macrophages were exposed to brain homogenates from tg338 mice infected with 127S and LA19K prions (1% (w/v) dilution in DMEM-10% FCS-PS) for 24h at 37°C. After two washes in D-PBS, the cells were lyzed or incubated for 9 days. At regular time points, the contents of the wells were lyzed in lysis buffer (0.5% sodium deoxycholate, 0.5 Triton X-100, 50 mM Tris-HCl, pH 7,4) and centrifuged for 1 min at 500 x g. The supernatants were collected and analyzed for protein content (MicroBCA kit, Pierce). The equivalent of 250 μg of proteins was digested by 1μg of PK for 1h at 37°C. The samples were then methanol precipitated, resuspended in Laemmli buffer, denatured and analyzed for PrP^res^ content by western blot.

## Results

### Distinct PrP^res^ types in the brain and spleen of high-expressor mice intracerebrally inoculated with LAN-like and CH1641-like sheep scrapie isolates

The tg338 mice overexpress the VRQ allele of ovine PrP. The PrP^C^ levels in the brain are ∼8-fold higher than in the sheep brain [29]. The spleen-to-brain PrP^C^ ratio is ∼1:20 in tg338, as in wild-type mice [22]. We transmitted by intracerebral (IC) route 37 sheep scrapie isolates from various countries and genotypes (including VRQ) to tg338 mice (details in supplemental Table 1). The isolates were from the LAN group [29] or closely resembling to CH1641 sheep scrapie isolate ([41–44], Figure S1). Sheep brains from the LAN group exhibit a PrP^res^ electrophoretic signature with unglycosylated fragments migrating around 21 kDa (21K-PrP^res^, Figure 1A). These cases are composed of two prion strain types termed LA21K and LA19K, which are preferentially selected in transgenic mice expressing low or high levels of PrP^C^, respectively [29]. The CH1641-like isolates exhibit a 19 kDa signature in the brain, as CH1641 (19K-PrP^res^, Figure 1A, Figure S1, [42, 43]). Initially, these isolates were found to exhibit a signature similar to BSE in sheep and were termed BSE-compatible isolates (Figure S1, [41]). However, transmission of CH1641-like isolates to ovine PrP mice (ARQ or VRQ allele) led to isolation of prions with strain features similar to LA19K prions ([41–43] and this study). For comparison, the PG127 sheep scrapie isolate (Figure 1A), which led to the accumulation of the same prion strain type in tg338 brain and spleen was used [45]. All the inoculated isolates induced a typical prion disease in tg338 mice. The PrP^res^ electrophoretic signature was compared between the brain and spleen of the same animals at the terminal stage of disease, so as to determine which substrain component was preferentially replicated. The results are summarized in Table 1 and a representative immunoblot is shown in Figure 1A.

**Figure 1.**
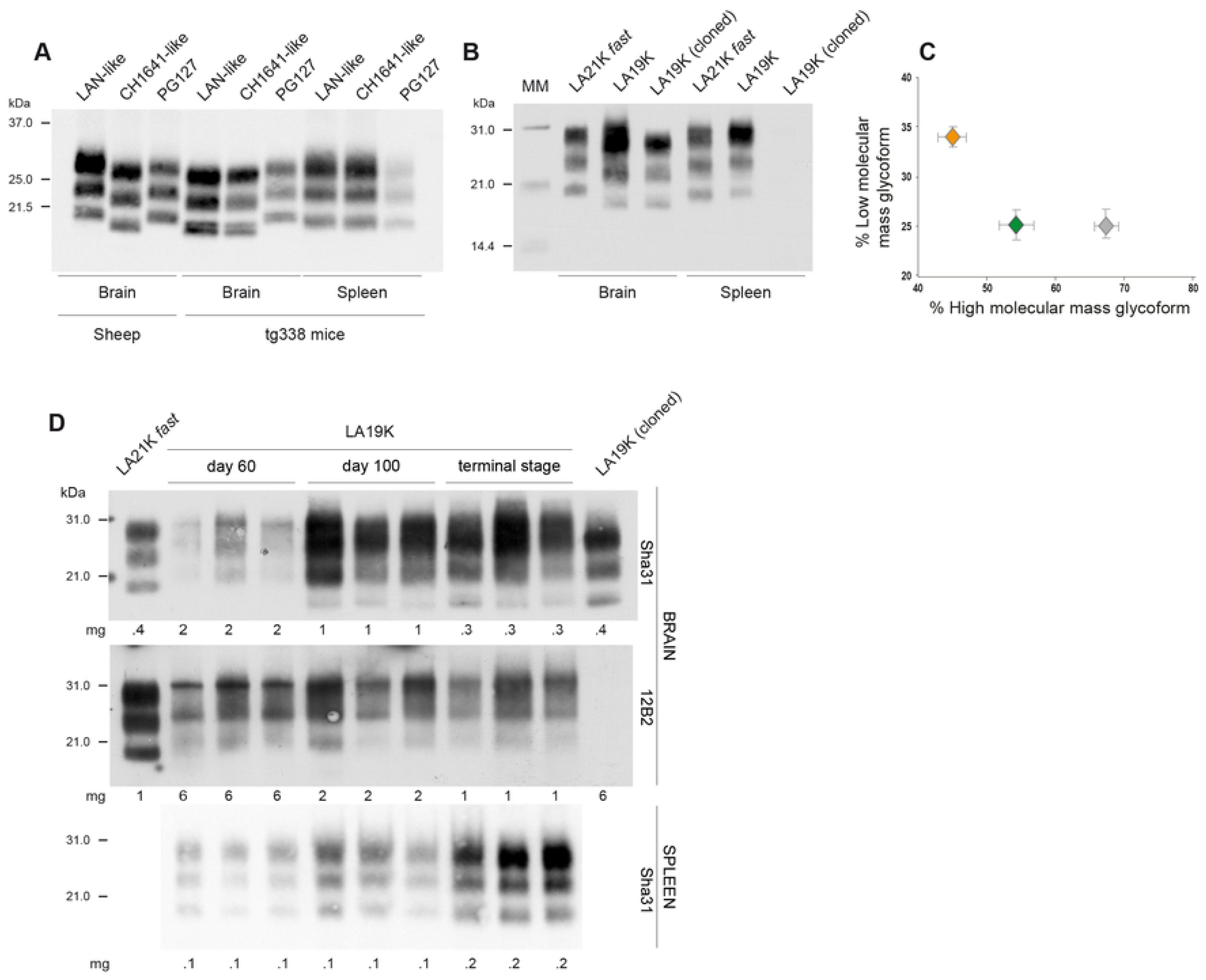
PrP^res^ electrophoretic pattern in brains and spleens of high-expressor tg338 mice inoculated IC with LAN-like and CH1641-like isolates. A) Distinct PrP^res^ electrophoretic pattern in the brain and spleen of tg338 high-expressor mice following IC challenge with representative LAN-like and CH1641-like isolates. The patterns observed after PG127 inoculation are shown as control. The PrP^res^ profile in the sheep brain are shown for comparison on the left side of the western blot. B) PrP^res^ detection and electrophoretic pattern in the brain and spleen of tg338 mice on serial transmission by IC route (6^th^ passage) of LAN404 isolate (LA19K), cloned LA19K and LA21K *fast* prions. MM: molecular mass markers. C) Ratio of diglycosylated *vs* monoglycosylated PrP^res^ in the spleen of mice on serial transmission of LAN404 isolate (gray symbol), LA21K *fast* (green symbol) and PG127 (orange symbol) (*n=5* spleens analyzed at the 6^th^ passage, data plotted as mean ±SEM). D) Time course analysis of PrP^res^ accumulation in the brain and spleen of tg338 mice on serial transmission (5^th^ passage) of LAN404 isolate (LA19K). Three mice were analyzed at each time point. The immunoblots were revealed with Sha31 and 12B2 anti-PrP antibodies, as indicated. PrP^res^ detection with these antibodies was also performed with tissues from terminally-sick mice after infection with cloned LA19K and LA21K *fast* prions. The amount of mg tissue loaded on the gels is indicated.

**Table 1.** PrP^res^ electrophoretic pattern in the brain and spleen after intracerebral inoculation of sheep scrapie field isolates to tg338 mice

| TSE sources | Number<sup>a</sup> | PrP<sup>res</sup> type<sup>b</sup> |  |  |
| --- | --- | --- | --- | --- |
|  |  | Sheep brain | tg338 brain (n/n<sub>0</sub>)<sup>c</sup> | tg338 spleen (n/n<sub>0</sub>)<sup>c</sup> |
| LAN-like | 26 | 21K | 19K (77/79) | 21K (53/53) |
| CH1641-like | 11 | 19K | 19K (38/38) | 21K (18/21)<sup>d</sup> |
| PG127 | 1 | 21K | 21K | 21K |

<sup>a</sup>Number of isolates transmitted to tg338 mice
<sup>b</sup>As referred to the size of unglycosylated PrP<sup>res</sup> in immunoblots
<sup>c</sup>Number of mice with the PrP<sup>res</sup> type/number of mice tested
<sup>d</sup>3 spleens were PrP<sup>res</sup> negative

After IC inoculation of LAN- and CH1641-like isolates, all but two tg338 brains analyzed showed prominent accumulation of 19K-PrP^res^, as reported previously with the prototypal LAN404 isolate [29]. The resulting agent was termed LA19K because of this prominent 19K-PrP^res^ signature in the brain [29]. In striking contrast, nearly all the spleens analyzed exhibited a 21K-PrP^res^ signature. The distinct PrP^res^ patterns in the brain and spleen from the same mice were observed over iterative passages of brain material by IC route, as illustrated with the LAN404 isolate (LA19K at the 6^th^ passage, Figure 1B).

Transmission of sheep scrapie isolates from the LAN group occasionally led to the emergence of a highly pathogenic ‘mutant’ designated LA21K *fast* in a minority of tg338 mice [29]. IC-inoculation of LAN-like isolates resulted in the emergence of LA21K *fast* in two out of the 79 brains analyzed (Table 1).

As LA21K *fast* prions accumulate in tg338 mouse spleens (Figure 1B and [46]), we questioned whether the 21K-PrP^res^ signature observed in the spleen on IC inoculation of LAN-like and CH1641-like isolates could be similar to the LA21K *fast* one. The strain-specific [47] glycoform profile from LA21K *fast* PrP^res^ in the spleen was distinct from that of the 21K-PrP^res^ found in the spleen on serial passage of LAN404, the latter being enriched in diglycosylated forms (Figure 1B-C). The prion types accumulating in the spleen on IC inoculation of LAN-like and CH1641-like isolates are thus likely different from LA21K *fast* prions.

A time-course analysis of PrP^res^ accumulation was performed in the brain and spleen tissue at the 5^th^ passage of LAN404 in tg338 mice. As shown in Figure 1D, 21K-PrP^res^ accumulation plateaued in the spleen from day 60 post-infection onward (bottom panel).

In the brain, 21K-PrP^res^ was detected at day 60. At day 100 and at terminal stage of the disease, 19K-PrP^res^ was mostly detected (top panel). The use of the 21K-selective anti-PrP monoclonal antibody 12B2 [38] to reveal the immunoblots demonstrated that 21K-PrP^res^ was still present in the brain at these time points (middle panel). This suggested co-propagation of the 19K-PrP^res^ and 21K-PrP^res^ components in the brain.

To summarize, while IC transmission of LAN-like and CH1641-like isolates to tg338 mice leads to the isolation of LA19K prions in the brain, a divergent, prominent 21K-PrP^res^ signature was observed in the spleen. This 21K-PrP^res^ signature, also subdominant in the brain, differed from that of LA21K *fast*, suggesting that LA21K prions replicated in the spleen of the infected mice.

### Co-propagation of LA19K and LA21K prion strain variants in the brain and spleen of high-expressor mice

To substantiate the view that the 21K-PrP^res^ signature in the spleen is associated with LA21K replication and formally exclude LA21K *fast* replication or a tissue-specific proteolytic processing of the LA19K strain type [48], we transmitted (IC route) spleen and brain extracts from the same tg338 mouse infected with LA19K (3^rd^ passage of LAN404) to reporter tg338 mice. We also challenged intracerebrally low-expressor tg143^+/-^ mice (expressing 1.5-fold PrP^C^ compared to sheep brain [29]), as they allow the dominant propagation of LA21K prions in the brain [29]. The resulting disease phenotypes in the two mouse lines are summarized in Figure 2A. In tg338 mice, the mean incubation duration (ID) was >3-fold longer with the spleen than with the brain extract. The spleen ID was also 3-fold longer than LA21K *fast* ID at the limiting dilution [6, 29]. The PrP^res^ pattern found after inoculation of spleen material was 21K in all the brains and spleens analyzed (Figure 2A-B). The neuroanatomical deposition pattern of PrP^res^ strikingly differed after inoculation of brain and spleen. In particular, numerous plaque-like PrP^res^ deposits were seen specifically after inoculation of the spleen extract (Figure 2C). This phenotype was reminiscent of LA21K prions [29]. Similar isolation of LA21K prions in tg338 mouse brains was observed on IC transmission of spleens from tg338 mice inoculated with other LAN- or CH1641-like isolates (Figure S2). In tg143^+/-^ mice, the spleen and brain extracts induced a long incubation time, with a 21K-PrP^res^ pattern. Such characteristics were reminiscent of LA21K prions in these mice (Figures 2A-B, [29]). To consolidate the view that the 19K-PrP^res^ and the 21K-PrP^res^ components were independent replicative entities in tg338 mice, we investigated the brain versus spleen colonization following IC challenge with cloned LA19K prions (double biological cloning by limiting dilution at the 3^rd^ iterative passage of LA404 [29]). 19K-PrP^res^ was detected in the brain (Figure 1B), with no evidence for the co-presence of a minor 21K-PrP^res^ signature, as assessed with 12B2 antibody (Figure 1D). All the spleens of the terminally-sick animals were PrP^res^-negative (Figure 1B-1D). Reporter tg338 mice were inoculated IC with these negative spleens to estimate the amount of LA19K infectivity by an incubation time bioassay. As shown in Figure 2A, the spleen extracts induced disease in 2 out of 8 mice >300 days post-infection, with a 19K-PrP^res^ signature in the brain. On the basis of such ID and the low attack rate [6], the spleen of biologically cloned LA19K prions harbored 10^5^-fold less infectivity than the brain.

**Figure 2.**
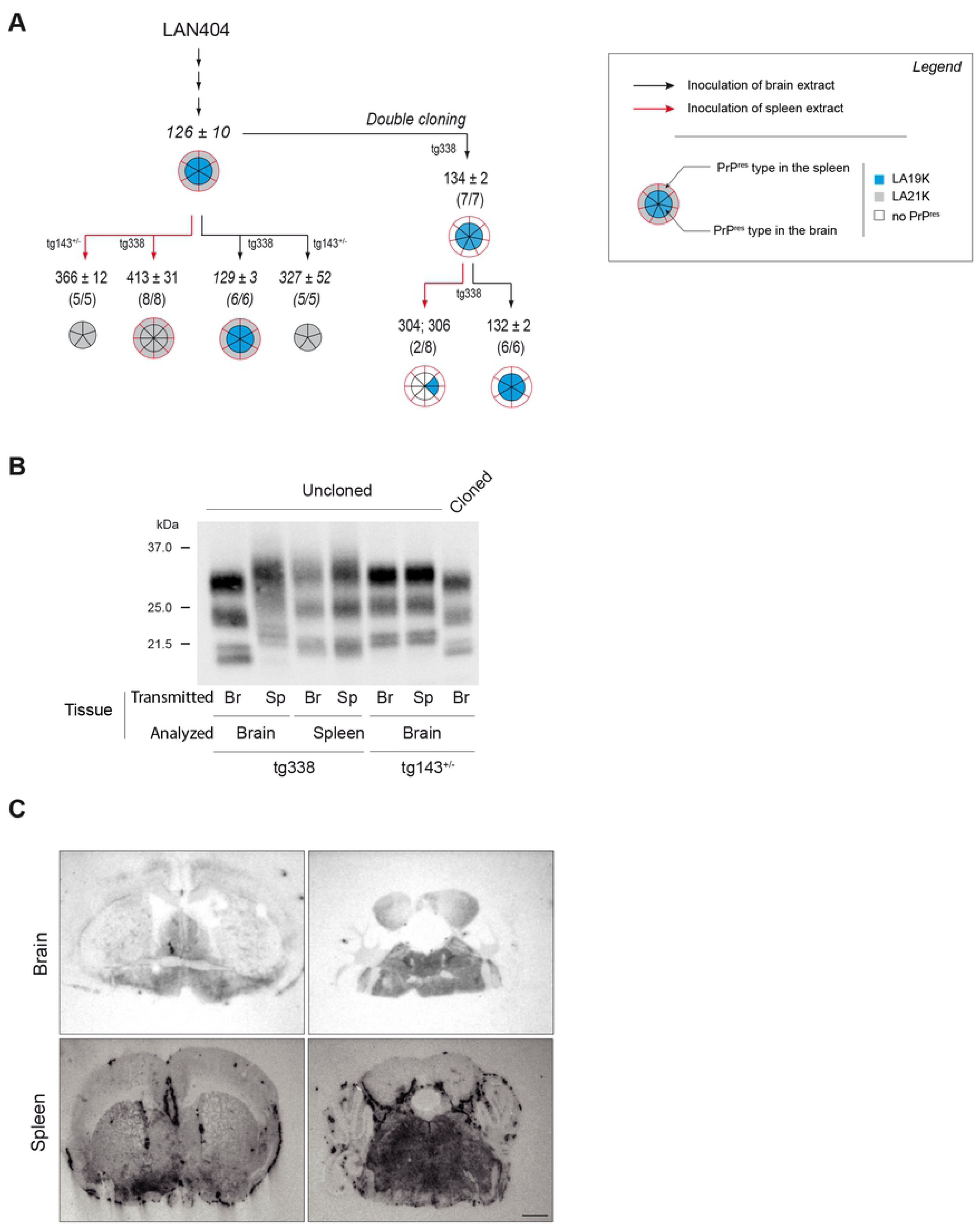
Strain phenotype of prions replicating in tg338 mouse spleens on serial passage of LAN404 isolate. A) Summary of the transmissions by IC route of brain or spleen extracts from tg338 mice infected with 19K prions (3^rd^ passage of LAN404) or with bicloned LA19K prions to reporter tg338 mice or to tg143^+/-^ mice expressing ovine PrP^C^ at near physiological levels. Transmission with brain or spleen extracts are indicated with black and red lines, respectively. The number of affected/inoculated mice (mice with TSE and positive for brain PrP^res^ by immunoblot) and the mean survival times in days ± SEM are indicated for each inoculated group. Segmented, doubled circles are used to indicate the proportion of mice with 19K PrP^res^ signature (blue), 21K PrP^res^ signature (grey) or absence of PrP^res^ in the brain (inside of the circle, black lines) and the spleen (outside of the circle, red lines). The data shown are representative of 5 independent transmission experiments with different mice infected with LA19K or cloned LA19K prions. Data in italic are from [29]. B) Representative immunoblot showing PrP^res^ electrophoretic pattern in brain and spleen from high-expressor tg338 mice and low-expressor tg143^+/-^ mice, depending on whether tg338-passaged brain (Br) or spleen (Sp) material (3^rd^ tg338-passage of LAN404) is used for IC inoculation. C) Representative histoblots at the level of the striatum (left panel) and the brain stem (right panel) showing the distribution of PrP^res^ deposits in the brains of high-expressor tg338 mice inoculated with brain or spleen extracts (3^rd^ tg338-passage of LAN404). Scale bar, 1 mm.

Collectively, these data indicate that IC inoculation of LAN- and CH1641-like isolates to tg338 mice results in the preferential replication of the LA19K and LA21K subcomponents in the brain and spleen, respectively. In the brain, both LA19K and LA21K subcomponents coexist, with the LA19K subcomponent in higher proportion. In the spleen, LA21K prions dominate, LA19K prions being barely detectable. LA19K prions thus show a preferential tropism for tg338 mouse brain. Such tissue-dependent segregation is reminiscent of the diverging selection of LA19K and LA21K prions on transmission to transgenic mice expressing PrP^C^ at varying levels [29].

### Expression of LA21K prions in high-expressor mouse brain on intraperitoneal inoculation of the LAN-like and CH1641-like isolates

Intraperitoneal (IP) inoculation primarily favors the neuroinvasion of prions that primo-replicate in the spleen, by transport through the peripheral nervous system and the spinal cord (review [49]). We thus asked whether extraneural inoculation of LAN-like and CH1641-like isolates would lead to the dominant expression of LA21K prions in the brain as LA21K prions rapidly colonized the spleen of high-expressor tg338 mice. A group of representative sheep scrapie sources was transmitted to tg338 mice by IP route. The dose injected was the same as for the IC route. Four isolates were from the LAN-like group (LAN404, ARQ16, 378, 454) and 2 from the CH1641-like group (O100 and m48). The PG127 isolate and LA19K prions served for comparison. Mice were euthanized at regular time-points post-injection and at terminal stage of disease (or end life) to examine early spleen colonization and analyze the molecular PrP^res^ profile in both the spleen and the brain. In conventional mouse models inoculated IP with splenotropic prions, PrP^res^ is detected at early time-points and the mean IDs are ∼1.3-fold prolonged compared to IC infections at similar dose [50–53]. Similar features were found with the PG127 isolate with regard to early PrP^res^ positivity in the spleen (from day 30) and ID prolongation (Figures 3A-B; supplemental Table S2). Similarly, the spleens were uniformly and early PrP^res^-positive following IP inoculation with the LAN-like and CH1641-like sheep scrapie isolates (Figures 3A-B). Yet, the tempo and the clinical manifestation of the disease and the dominant strain type replicating in the brain were not uniform amongst the mice analyzed. The mean IDs were ∼3-fold longer than in the IC inoculated animals, ranging from 520 to 560 days (Figure 3A). Twenty-four percent of the brains analyzed were PrP^res^-negative. Fifty-nine percent of the brains exhibited a 21K-PrP^res^ pattern and sixteen percent a 19K-PrP^res^ pattern (Figures 3A-B). The 21K-PrP^res^ pattern dominated over the 19K-PrP^res^ pattern for all but one isolate (ARQ16). We conclude that IP inoculation of LAN-like and CH1641-like isolates favors the neuropathological expression of the splenotropic LA21K prions at the expense of LA19K prions. The significant proportion of mice with no PrP^res^ in the brain despite early, widespread accumulation of PrP^res^ in the spleen suggests a delayed efficacy of LA21K prions to replicate at full attack rate in the brain following IP inoculation.

**Figure 3.**
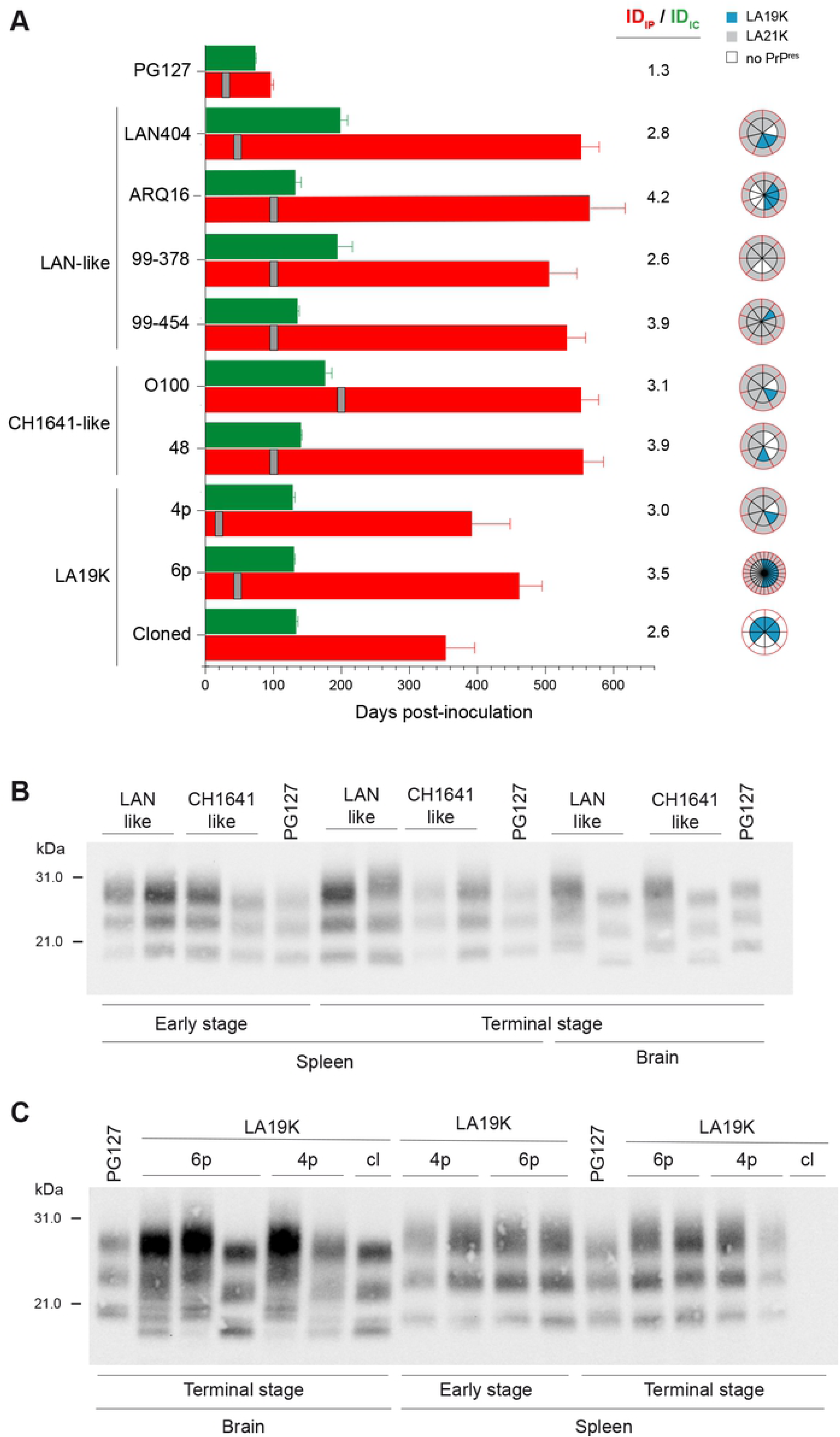
PrP^res^ electrophoretic pattern in brains and spleens of high-expressor tg338 mice inoculated IP with LAN-like and CH1641-like isolates. A) Comparison of the transmissions by IP versus IC route of PG127, LAN-like, CH1641-like sheep scrapie isolates and LA19K prions (4^th^ and 6^th^ IC-brain-passage (4p, 6p) and cloned agent). The mean ±SEM incubation durations (ID) are indicated following challenge by IC route (green bar) and IP route (red bar). The ID_IP_ / ID_IC_ ratio are indicated. The grey vertical bar indicates the time at which PrP^res^ was first detected in the spleen (Supplemental Table S2). Segmented, doubled circles are used to indicate the proportion of mice with 19K PrP^res^ signature (blue), 21K PrP^res^ signature (grey) or absence of PrP^res^ in the brain (inside of the circle, black lines) and the spleen (outside of the circle, red lines). B) Representative immunoblot showing PrP^res^ electrophoretic pattern in brain and spleen from tg338 high-expressor mice inoculated IP with isolates from the LAN-like and CH1641-like groups, at the terminal stage of disease. The spleen and the brain from the same animal are presented, meaning that the brain can harbor a 21K or 19K PrP^res^ signature whereas a unique 21K signature is detected in the spleen. The spleen PrP^res^ profiles are also shown at early time points post-inoculation (from 50 to 200 days depending on the isolate, Supplemental Table S2). Patterns after PG127 infection are shown for comparison. C) Representative immunoblot showing PrP^res^ electrophoretic pattern in brain and spleen from tg338 high-expressor mice inoculated IP with LA19K prions (4^th^ IC-brain-passage (4p), 6^th^ IC-brain-passage (6p) and cloned (cl)), at the terminal stage of disease. The spleen and the brain from the same animal are presented, meaning that the brain can harbor a 21K or 19K PrP^res^ signature (with the 6^th^ pass, a double 21/19K signature was observed) whereas a unique 21K signature is detected in the spleen The spleen PrP^res^ profiles are also shown at early time points post-inoculation (from 20 to 50 days depending on the number of passages, Supplemental Table S2). Patterns after PG127 infection are shown for comparison.

We studied whether changing the inoculation route similarly impacted the pathogenesis of LA19K prions. Two distinct passages were used, the 4^th^ and the 6^th^ IC-brain passage of LAN404. We detected an early accumulation of 21K-PrP^res^ in all the spleen analyzed, independently of the number of passages (Figures 3A, 3C). In the brain, the proportion of 21K-PrP^res^ mice decreased with the number of passages, at the expense of the 19K-signature. 71% and ∼45% of the brains analyzed were 21K-positive at the 4^th^ and 6^th^ passage of LAN404, respectively (Figures 3A, 3C). These results were consistent with an enrichment in LA19K prions over serial IC passaging.

IP inoculation of bi-cloned LA19K prions resulted in a disease at near full attack-rate, with 19K-PrP^res^ accumulating in the positive brains, whereas all the spleens remained PrP^res^-negative (Figures 3A, 3C). These findings suggest that LA19K prions could directly neuroinvade from the peripheral sites of infection, a pathway previously observed for other prions in situation of impaired replication in the spleen [20, 21, 54]. However, the ID was still ∼2.6-fold longer than by IC route, approaching values obtained with primary sheep material or uncloned LA19K prions (Figure 3A), indicating a delayed efficacy of LA19K prions to neuroinvade from the periphery.

### Preferential selection of LA21K prions on inoculation of high-expressor mice with spleen material from LAN404 primary isolate

We asked whether a similar distinctive splenotropism between LA21K and LA19K prions pre-existed in the original sheep isolate. A spleen extract from the LAN404 isolate was inoculated by IC and IP routes to tg338 mice. At variance with the IC-inoculation of LAN404 brain material which resulted in dominant expression of LA19K prions in the brain and LA21K in the spleen ([29] and Figure 4A), IC-inoculation of LAN404 spleen material led to accumulation of prions with a 21K-PrP^res^ signature in both brain and spleen in ∼450 days (Figure 4A-B). After IP inoculation, the incubation time established at 660 days and all the brains and spleens analyzed exhibited a 21K-PrP^res^ signature (Figure 3A-B).

**Figure 4.**
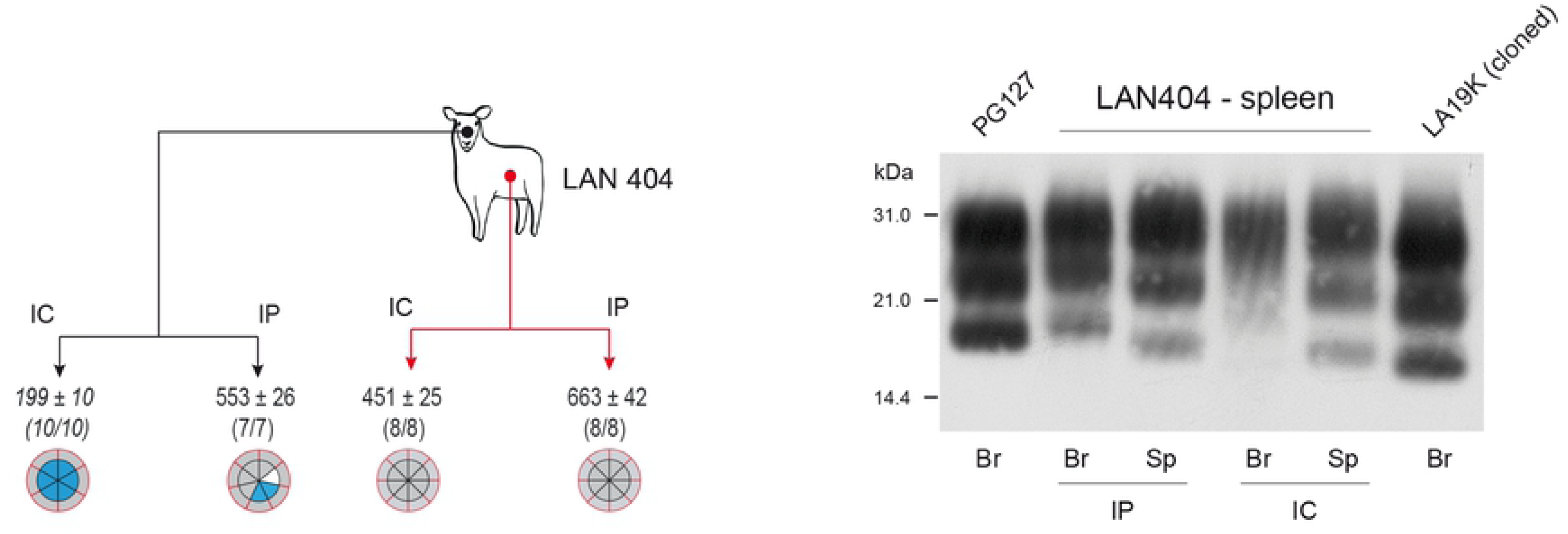
Strain phenotype of prions replicating in tg338 mouse spleen and brain on primary passage of LAN404 spleen. A) Summary of the transmissions by IC or IP route of brain or spleen extracts from LAN404 sheep scrapie isolate to tg338 mice. Legends are as in Figure 2. Data in italic are from [29]. B) Representative immunoblot showing PrP^res^ electrophoretic pattern in (Br) and spleen (Sp) from high-expressor tg338 mice, upon inoculation with LAN404 spleen material by IP or IC route. PrP^res^ electrophoretic pattern of LA21K *fast* and cloned LA19K in tg338 mouse brain is shown for comparison.

We conclude that in the spleen of the natural host, the LA21K component predominates over the LA19K component to levels allowing molecular expression in the CNS and in the spleen on primary transmission to high-expressor mice.

### Another pair of strains co-propagating in high-expressor mice and with differing tropism for the brain and spleen shows divergent selection according to PrP^C^ expression level

To further explore the possibility that prion capacity to replicate in the spleen is controlled by PrP^C^ expression levels, we compared the replicative capacity in mice expressing low levels of PrP^C^ of two pairs of strains with opposite splenotropism in high expressors. Previously, we reported that a pair of prion strains, designated T1^Ov^ and T2^Ov^, isolated through the adaptation of cortical MM2 sporadic CJD (MM2-sCJD) prions to tg338 mice exhibited differing tissue tropism. T2^Ov^ replicated dominantly in the brain with T1^Ov^ being present as a subcomponent, while T1^Ov^ preferentially populated the spleen ([28] and Figure 5A-B). A brain extract from tg338-passaged MM2-sCJD prions (4^th^ passage) was inoculated by IC route to tg335^+/-^ mice, expressing 1.2-fold PrP^C^ compared to sheep brain [29]. For comparison, brain extracts containing solely T2^Ov^ prions (obtained by bicloning by limiting dilution [28]) or T1^Ov^ prions (PMCA selection and single cloning [28]) were inoculated to tg335^+/-^ mice. The transmission of tg338-passaged MM2-sCJD prions to tg335^+/-^ mice led to the dominant expression of T1^Ov^ prions, as based on the mean ID (Figure 5A), the PrP^res^ signature in brain and spleen by western blot (Figure 5B) and the neuroanatomical distribution of PrP^res^ by histoblotting (Figure 5C), which all showed the T1^Ov^ characteristics observed after infection with cloned T1^Ov^ prions. The apparent counter-selection of T2^Ov^ prions in tg335^+/-^ mice was not due to their incapacity to replicate on low-expressor mice as bicloned T2^Ov^ prions elicited disease in these mice in ∼330 days (Figure 5A) with a T2^Ov^-PrP^res^ specific banding pattern in the brain (Figure 5B) and presence of low levels of PrP^res^ by histoblotting (Figure 5C). Together, these data indicate that for another pair of co-propagating prion strains, there is a tight correlation between capacity to replicate in the spleen and capacity to be dominantly expressed on mice expressing low PrP^C^-levels in the brain.

**Figure 5.**
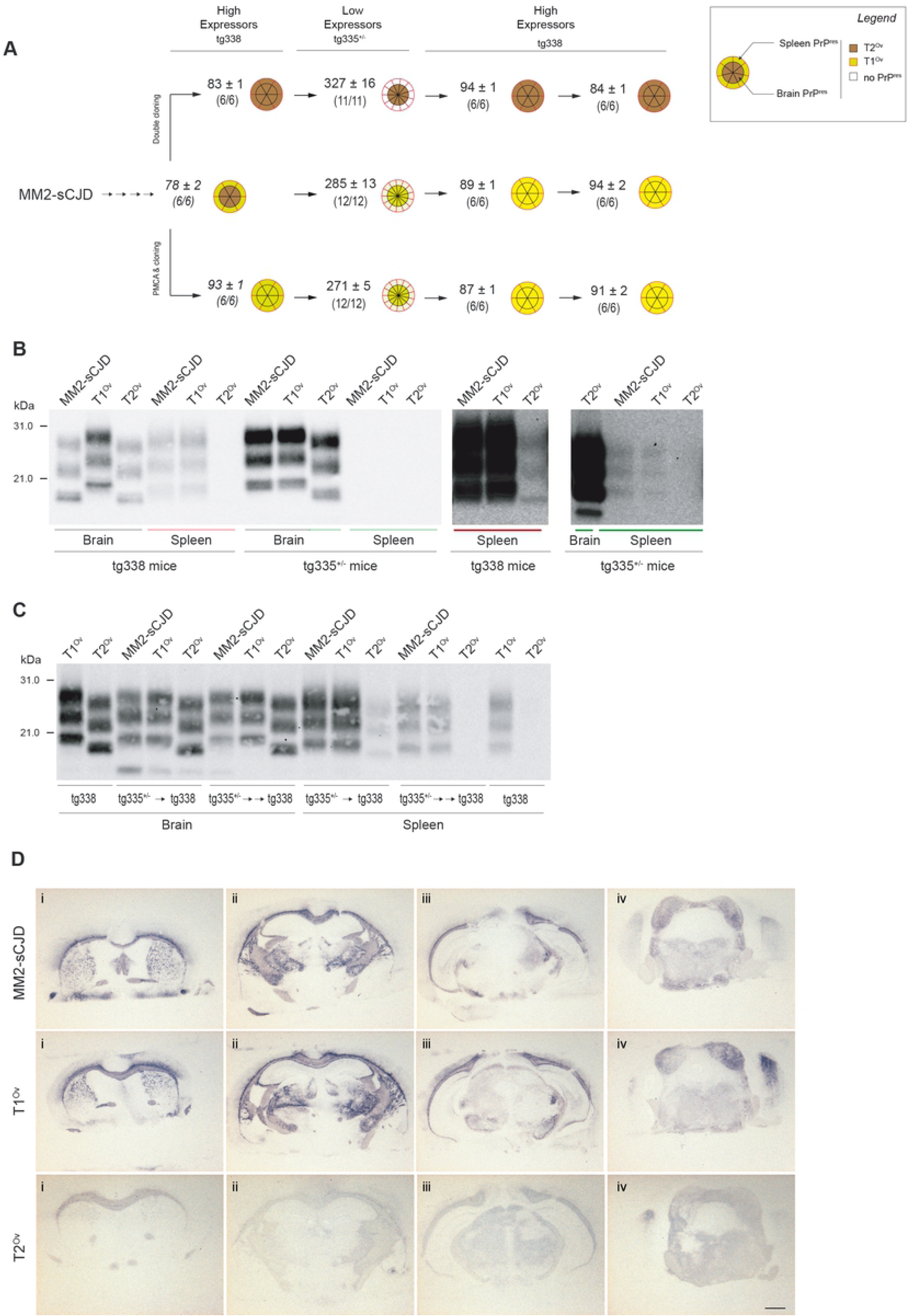
PrP^C^-level dependent selection of prions with differing splenotropism in ovine PrP transgenic mice. A) Summary of the transmissions by IC route of brain extracts from tg338 mice infected with MM2-sCJD prions (4^th^ passage) or with cloned T1^Ov^ and T2^Ov^ prions to tg335^+/-^ mice expressing ovine PrP^C^ at near physiological levels, before back passage to tg338 mice. The number of affected/inoculated mice (mice with TSE and positive for brain PrP^res^ by immunoblot) and the mean survival time in days ± SEM are indicated for each inoculated group. Segmented, doubled circles are used to indicate the proportion of mice with T2^Ov^ (brown or clear brown depending on the accumulation levels) or T1^Ov^ (yellow or clear yellow) PrP^res^ signatures in their brains (inside of the circle, black lines) and spleens (outside of the circle, red lines). Data in italic are from [28]. B) Representative immunoblot showing PrP^res^ electrophoretic pattern in brain and spleen from tg338 high-expressor mice and tg335^+/-^ low-expressor mice infected with MM2-sCJD prions (4^th^ passage) or with cloned T1^Ov^ and T2^Ov^ prions. The gels on the right side are portions of the gels (colored red and green lines) that are purposely overexposed to detect T2^Ov^ PrP^res^ in the spleen. C) Representative immunoblot showing PrP^res^ electrophoretic pattern in brain and spleen from tg338 high-expressor mice after one or two back passages from tg335^+/-^ low-expressor mice infected with MM2-sCJD prions or with cloned T1^Ov^ and T2^Ov^ prions. PrP^res^ signatures from T1^Ov^ and T2^Ov^ prions in tg338 mice are shown for comparison. D) PrP^res^ deposition pattern in the brain of tg335^+/-^ low-expressor mice infected with tg338-passaged MM2-sCJD, T1^Ov^ and T2^Ov^ prions. Representative histoblots of antero-posterior coronal brain sections at the level of the septum (i), hippocampus (ii), midbrain (iii) and brainstem (iv). Scale bar, 1 mm.

Remarkably, the dominant expression of T1^Ov^ prions on passage of tg338-passaged MM2-sCJD prions to tg335^+/-^ mice occurred to the detriment of T2^Ov^ prions. Over two serial back passages to tg338 mice, T2^Ov^ prions were not phenotypically recovered. The T1^Ov^ signature remained dominant in all brains and spleens from tg338 mice (Figure 5A, 5C). Such outcome was not observed on retro passage of LAN-like isolates from low- to high-expressor mice as LA19K prions were recovered after intermediate passage onto low-expressors [29]. Alternance of transmission in mice expressing variable levels of PrP^C^ may thus result in a strain evolution as strong as that observed in a heterotypic PrP transmission context.

## Discussion

Building on our previous observations that the phenotypic expression of two prion subcomponents present in variable proportions in numerous sheep scrapie isolates is driven by PrP^C^ expression levels in transgenic mice, we now show that the segregation between these two agents occurs between the brain and the spleen of the same infected high-expressor mouse, these tissues expressing different PrP^C^ levels. Further, the replicative dominance in the brain of a pair of strains derived from sporadic CJD prions with opposite tropism for the brain and spleen appears tightly controlled by PrP^C^ expression levels in transgenic mice expressing varying PrP^C^ levels. Host PrP^C^ levels may thus impact prion (in)capacity to replicate in the lymphoid tissue.

LA19K prions predominated in high-PrP^C^ expressor tg338 mouse brains inoculated IC with a large panel of natural sheep scrapie isolates composed in variable proportion of LA19K and LA21K prions (LAN-like, [29] and this study). Here, we show early, widespread and dominant accumulation of LA21K in the spleen of these mice, -as based on molecular strain typing, and phenotypic characterization on high- and low-PrP^C^ expressor mice. The LA19K/LA21K segregation was also observed after IC inoculation of CH1641-like isolates in which the 19K subcomponent was originally dominant, indicating that the lymphoid tissue is a privileged site for LA21K prions replication. In contrast, cloned LA19K prions replicated with limited efficacy in the spleen, and thus were considered as mostly neurotropic.

That co-propagating TSE substrains replicate preferentially in distinct tissues from the same host is not restricted to transgenic mice nor to the VRQ allele of ovine PrP. We show here a similar enrichment of LA21K prions in the spleen relative to brain in the LAN-infected sheep. A similarly distinct selection of PrP^res^ conformers with 19K and 21K signature was observed in the brain and spleen tissue from transgenic mice expressing the ARQ allele of ovine PrP inoculated with CH1641-like scrapie isolates [55]. A similar tissue-specific segregation of prion subcomponents due to distinct tropism was observed on transmission of a human vCJD case to transgenic mice overexpressing human PrP [23, 56], on transmission of cortical MM2-sporadic CJD to tg338 mice [28], directly in a sporadic CJD individual [57] and in CWD-infected elk [58].

These data advocate for strain typing in both brain and spleen tissues, including in high expressor mice. Blind strain typing of LAN-like isolates using tg338 mouse brain only would categorize them as CH1641 prions. However, examining the lymphoid tissue or changing the route of infection allows expression of the LA21K strain component, consistent with studies in transgenic mice expressing variable PrP levels. *In fine* transgenic mice in which PrP is introduced by additional transgenesis prove to be a valuable tool to reveal natural prion strain diversity present in TSE isolates, provided the PrP-encoding constructs allow correct PrP^C^ expression in the lymphoid tissue.

The host and structural requirements for prion replication in the lymphoid tissue remain mostly unknown. From the prion pathogen viewpoint, two broad hypotheses recently emerged, the sensitivity to degradation of the PrP^Sc^ assemblies and their aggregation size. A study from Shikiya et al. suggested that absence of DY prions replication in hamster spleens was due to a particular sensitivity of DY PrP^Sc^ to protease digestion [59], yet without providing *in vivo* or *ex vivo* evidence (e.g., clearance of PrP^Sc^ by macrophages). Here, we firmly exclude this possibility as LA19K prions were not more sensitive than 127S prions to either PK digestion or clearance by peritoneal macrophages (Figure 6). Aguilar-Calvo et al. proposed that prion conformations could impact pathogenesis after peripheral infection, i.e., highly fibrillar prion assemblies are poor neuroinvaders compared to subfibrillar prions but high replicators in the spleen [31]. PrP^Sc^ assemblies’ particulate profile may indeed impact access to the FDC from the site of infection [60, 61]. LA21K prions and vCJD prions would follow this rule as they would be categorized as ‘highly fibrillar’, as based on their sedimentation velocity profile and the presence of plaque-like deposits in the brain (this study and [23, 62]). However, T1^Ov^ and T2^Ov^ prions would escape such a rule as they exhibit similar ‘subfibrillar’ aggregation size [63], despite distinct tropism for the spleen.

**Figure 6.**
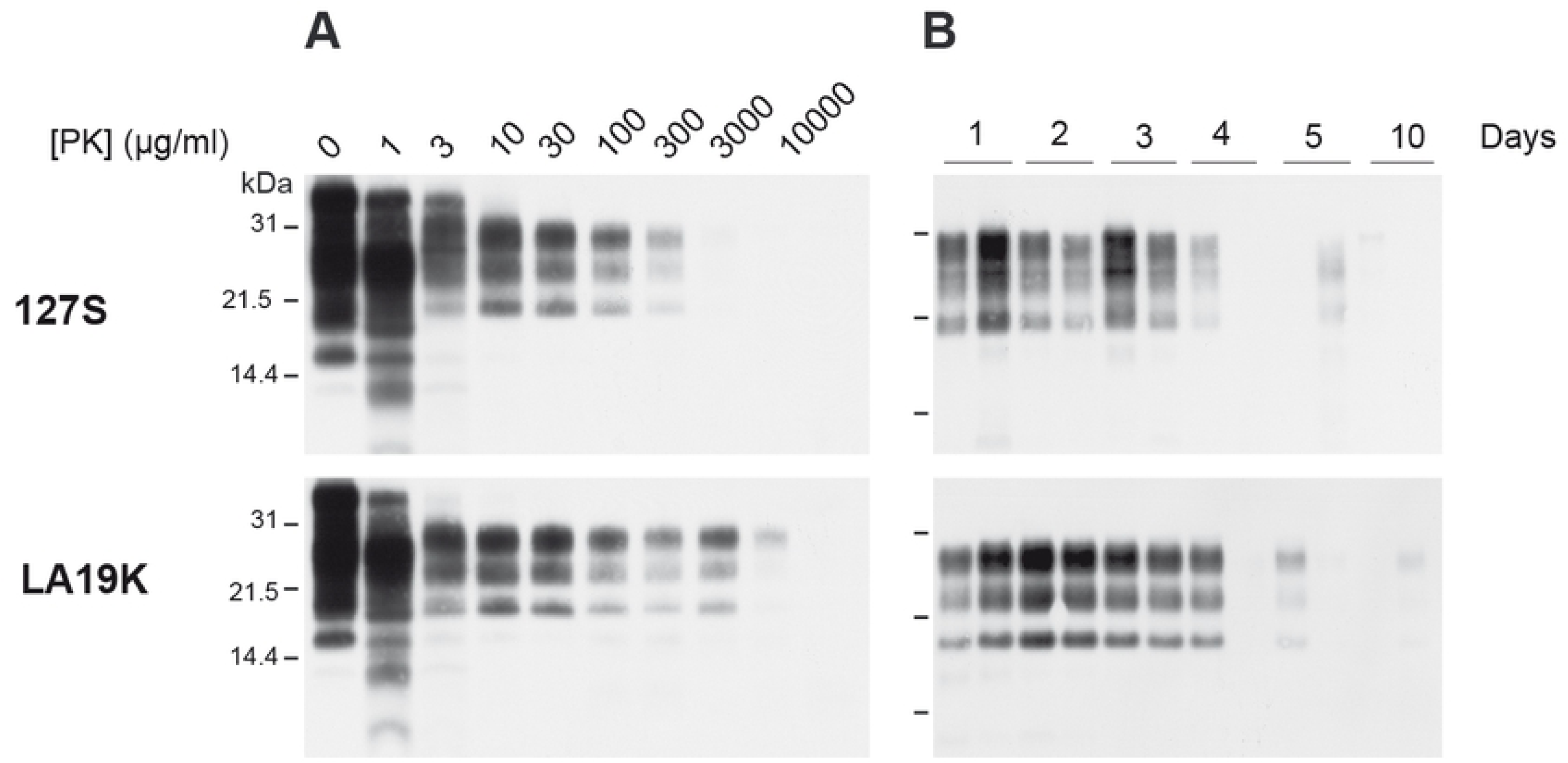
PrP^res^ electrophoretic pattern in brains and spleens of high-expressor tg338 mice inoculated IC with LAN and CH1641-like isolates. A) Comparative PK digestion assay performed on brain material from tg338 inoculated with 127S or cloned LA19K prions. The homogenates were treated with increasing concentrations of PK, as indicated, and analyzed by western blotting. B) Representative immunoblot showing PrP^res^ degradation in tg338 peritoneal macrophages exposed to 127S and LA19K prion for the indicated period of time (duplicates analysis).

From the host viewpoint, mature FDC are necessary [9, 15–21, 64, 65], being ontogenetically present or induced by chronic inflammation [66]. Co-factors have been proposed to be involved, including notably complement [67, 68], but the differential requirement amongst strains is not known. Here, LA19K / LA21K prions preferential targeting for brain and spleen correlated with PrP^C^ expression levels in these tissues. The superimposable PrP^C^-dependent selection obtained with a second pair of prion strains isolated from MM2-sCJD in tg338 mice and showing differing tropism for the lymphoid versus the cerebral tissue, that is the most lymphotropic is preferentially replicated on low-PrP^C^ expressor mice and the most neurotropic on high-PrP^C^ expressor mice, further argues that PrP^C^ expression levels may be a major driver of prion (sub)strain selection in different tissues of the infected host.

Additionally, at the molecular level, PrP^C^ conformational landscape is likely to differ between spleen and brain, due to different trimming [48], glycosylation [69] including sialylation [70], and thus may impact conversion by certain PrP^Sc^ subspecies, as observed during cross-species prion transmission. A combination of both PrP^C^ expression levels and variations in spleen PrP^C^ conformations may well explain why intermediate passage of tg338-passaged-MM2-sCJD prions on low expressors resulted in the abrupt exclusion of T2^Ov^ prions on back passage to tg338 mice, an outcome that was reminiscent of that observed during cross-species transmission events (for reviews [7, 8]). Such factors may be drivers of within-host prion diversification/evolution during homotypic transmissions.

It may be noted that PrP^C^ expression levels are classically quantified in the whole spleen and in the whole brain, yet meaning that different variations in PrP^C^ levels could occur locally in prion-replicating cells [71]. The exact contribution of FDC to spleen PrP^C^ expression level is unknown due to the fact that these cells only account for 0.1-1% of the cells, making isolation and quantification of their PrP^C^ expression levels a highly challenging issue. However, we previously reported by immunofluorescence that PrP^C^ expression levels appeared low in tg338 FDC compared to neurons [22].

As a tentative explanation for strain-dependent polymerization rate as a function of PrP^C^ levels, we inferred by mathematical modelling that differences in the number of PrP^C^ monomers integrated in the growing PrP^Sc^ assemblies (different kinetic order of the templating process) could explain PrP^C^-dependent substrain selection [29]. LA19K growth would necessitate integration of more PrP^C^ molecules than for LA21K prions. In theory, the strain necessitating more PrP^C^ to replicate should also start accumulating in the spleen. However, the plateau of PrP^Sc^ accumulation observed in the spleen quite early after infection [10, 23, 45], possibly due to a cessation of replication activity [72, 73] would prevent the chance that the kinetically-disadvantaged substrain replicates in this tissue.

The IP route of inoculation, known to favor neuroinvasion of prions from the lymphoid tissue (review [49]) allowed dominant accumulation of LA21K PrP^res^ in the brain of a proportion of high-expressor mice at the expense of LA19K PrP^res^. The incubation time lengthening when IP and IC routes of infection were compared for LAN-like isolates was overtly prolonged as compared to PG127 in tg338 mice or other experimental models and a number of IP-inoculated tg338 mice remained PrP^res^-negative in the brain while their spleen was strongly and early positive. This suggests an impaired capacity of LA21K prions to establish at full attack rate in the brain, as previously observed with other prion types [23, 24]. Bicloned LA19K prions were able to establish in the brain after IP infection, despite absence of replication in the spleen, by a direct neuroinvasion process. However, direct neuroinvasion after IP infection usually results in ID similar to those observed after primo-replication in the spleen [20, 21]. This suggests that the IC versus IP ID was also aberrantly prolonged for bicloned LA19K prions. Comparison of ID on low-versus high-expressors after inoculation of LA21K *fast* versus LA19K prions show ‘abnormal’ prolongation of ID for the latter [29]. It is therefore likely that PrP^C^-limiting events are similarly occurring in high-expressor mice during prion neuroinvasion after IP inoculation. Of note, when LAN404 sheep spleen instead of brain was transmitted to tg338 mice, the IP versus IC injection delay observed (x1.47, Figure 4) was rather consistent with the ∼1.3 value found in other experimental models [50–53] and with PG127 in tg338 mice (Figure 3). As the sheep spleen is less enriched than the brain in LA19K prions, these data would collectively suggest that LA19K prions give the disease tempo after IP infection even if LA21K PrP^res^ molecular signature prions can be dominant.

In conclusion, these data and our previous report [29] suggest an emerging contribution of PrP as a molecular driver/modulator of within-host prion selection. Events similar to the brain/spleen segregation observed here could also occur directly within the brain owing to the regional heterogeneity in PrP^C^ levels and conformations in this tissue [74]. From a diagnostic point of view, given the apparent complexity of natural animal and human TSE isolates in terms of substrain heterogeneity [27, 75, 76], these data advocate to type extraneural tissues or fluids for a comprehensive identification of the TSE agents circulating in susceptible mammals.

## Acknowledgements

We thank the staff of Animalerie Rongeurs (IERP, INRA, 2018. Infectiology of fishes and rodent facility, doi: 10.15454/1.5572427140471238E12, Jouy-en-Josas, France) for animal care, N. Moriceau for excellent technical help, P. Clayette (Bertin Pharma, Montigny-le-Bretonneux, France) and J. Langeveld (Wageningen University, Lelystad, The Netherlands) for the kind gift of Sha31 and 12B2 anti-PrP monoclonal antibodies, respectively. We thank our colleagues for kindly providing sheep TSE sources.

## Data Availability

All relevant data are within the manuscript and its supporting information files. Data are fully available without restriction.

## Supporting Information Legends

**Figure S1. PrP^res^ electrophoretic profile from CH1641-like isolates** Electrophoretic pattern of PrP^res^ in the brain from CH1641-like cases, compared with CH1641 isolate [36], PG127 isolate [35] and sheep experimentally inoculated with BSE prions [77].

**Figure S2. Strain phenotype of prions replicating in tg338 mouse spleens on serial passage of LAN-like or CH1641-like isolates**. Transmissions by IC route of brain or spleen extracts from tg338 mice infected with the LAN-like isolate (99-378 isolate, 2^nd^ passage) or the CH1641-like isolate (O100 isolate, 2^nd^ passage) to reporter tg338 mice. Transmission with brain or spleen extracts are indicated with black and red lines, respectively. The number of affected/inoculated mice (mice with TSE and positive for brain PrP^res^ by immunoblot) and the mean survival times in days ± SEM are indicated for each inoculated group. Segmented, doubled circles are used to indicate the proportion of mice with 19K PrP^res^ signature (blue) or 21K PrP^res^ signature (grey)in the brain (inside of the circle, black lines) and the spleen (outside of the circle, red lines).

**Supplementary Table 1** TSE sources transmitted to tg338 mice.

**Supplementary Table 2** PrP^res^ signature in the brain and spleen after intraperitoneal inoculation of LAN, CH1641-like isolates and LA19K prions to tg338 mice.

